# Microbiota mediated plasticity promotes thermal adaptation in *Nematostella vectensis*

**DOI:** 10.1101/2021.10.18.464790

**Authors:** Laura Baldassarre, Hua Ying, Adam Reitzel, Sebastian Fraune

## Abstract

At the current rate of climate change, it is unlikely that multicellular organisms will be able to adapt to changing environmental conditions through genetic recombination and natural selection alo. Thus, it is critical to understand alternative mechanisms that allow organisms to cope with rapid environmental changes. Here, we used the sea anemone *Nematostella vectensis* as model to investigate the microbiota as putative source of rapid adaptation. Living in estuarine ecosystems, highly variable aquatic environments, *N. vectensis* has evolved the capability of surviving in a wide range of temperatures and salinities. In a long-term experiment, we acclimated polyps of *Nematostella* to low (15°C), medium (20°C) and high (25°C) temperatures, in order to test the impact of microbiota-mediated plasticity on animal acclimation. Using the same animal clonal line, propagated from a single polyp, allowed us to eliminate effects of the host genotype. Interestingly, the higher thermal tolerance of animals acclimated to high temperature, could be transferred to non-acclimated animals through microbiota transplantation. In addition, offspring survival was highest from mothers acclimated to high temperature, indicating the transmission of thermal resistance to the next generation. Microbial community analyses of the F1 generation revealed the transmission of the acclimated microbiota to the next generation. These results indicate that microbiota plasticity can contribute to animal thermal acclimation and its transmission to the next generation may represent a rapid mechanism for thermal adaptation.

## Introduction

Changes in the climate are proceeding worldwide at a rate never registered before and temperatures will rise dramatically in the coming decades. Species able to migrate could move toward new-favourable areas, but those that have limited dispersal capacities or are sessile will have only two options: adaptation or extinction. Traditional theory and research since the Modern Synthesis have focused on the balance of mutation and selection as the central explanation for the adaptation of populations to their environment and as the generator of phenotypic novelty. However, some organisms also have a remarkable ability to acclimate to environmental change during their lifetime.

The mechanisms for acclimation are generally assumed to be due to shifts in gene expression regulation ^1,2^. A focus on this factor alone is surely incomplete because the phenotype of an animal cannot be explained entirely by its genes. In 1927, the microbiologist Ivan E. Wallin hypothesized in his book, “*Symbionticism and the Origin of Species”*, that the acquisition of bacterial endosymbionts favours the origin of new species ^3^. Unlike the genes and regulatory regions of the genome, microbial composition can be rapidly modified by environmental cues, and may thus represent a mechanism for rapid acclimation and adaptions of individuals to a changing environment ^4–7^. Recently, the microbiota-mediated transgenerational acclimatization (MMTA) concept was proposed ^8^, suggesting that changes in microbiota assemblages, occurring in acclimating animals, may be passed on through generations to confer long-lasting resistance to changing environments by individuals and populations.

To be able to disentangle host genetic and microbial contributions to thermal acclimation, we took advantage of the model system *Nematostella vectensis* ^9^. *N. vectensis*, an anthozoan cnidarian, is a sedentary predator that resides exclusively in estuaries and brackish water environments, where it lives borrowed in sediments ^10^. It is a wide-spread species that has been found in both Pacific and Atlantic coasts of the US and of the UK. In their natural habitats, wild populations of *N. vectensis* experience high variations of salinity, temperature and pollutants ^11–16^. Under lab conditions, all the developmental stages are procurable on a weekly basis and spawning is induced by a shift in temperature and exposure to light ^17^. *N. vectensis* can be easily cultured in high numbers ^13^ and clonally propagated to eliminate genetic confounding effects. A detailed analyses of its microbiota revealed that *N. vectensis* harbors a specific microbiota whose composition changes in response to different environmental conditions and among geographic locations ^18^. Recently, has been shown that female and male polyps transmit different bacterial species to the offspring and that further symbionts are acquired from the environment during development ^19^. Furthermore, a protocol based on antibiotic-treatment was established to generate germ-free animals that allow controlled recolonization experiments to be conducted ^20^. All together, these characteristics make the sea anemone *N. vectensis* a uniquely informative model organism to investigate the effects of bacterial plasticity on thermal acclimation ^5^.

Here we used a clonal line of *N. vectensis* to characterise physiological and microbial plasticity of the holobiont under different thermal conditions, while eliminating the variability due to different host genotypes. Using microbial transplantations to non-acclimated polyps, we proved the ability of acclimated microbes to confer resistance to thermal stress. We further showed that thermal resistance to heat stress is transmitted to the next generation.

Altogether, we provide strong evidences that microbiota-mediated plasticity contributes to the adaptability of *N. vectensis* to high temperature and that the transmission of acclimated microbiotas represents a mechanism for rapid adaptation.

## Materials and methods

### Animal culture

All experiments were carried out with polyps of *N. vectensis* (Stephenson 1935). The adult animals of the laboratory culture were F1 offspring of CH2XCH6 individuals collected from the Rhode River in Maryland, USA ^13,17^ They were kept under constant, artificial conditions without substrate or light in plastic boxes filled with 1L ca. *Nematostella* Medium (NM), which was adjusted to 16‰ salinity with Red Sea Salt^®^ and Millipore H_2_O. Polyps were fed 2 times a week with first instar nauplius larvae of *Artemia salina* as prey (Ocean Nutrition Micro *Artemia* Cysts 430 - 500 gr, Coralsands, Wiesbaden, Germany) and washed once a week with media pre-incubated at the relative culture temperatures.

### Animal acclimation

A single female polyp from the standard laboratory culture conditions (16‰ ppt, 20°C) was isolated and propagated via clonal reproduction. When a total of 150 new clones was reached, they were split into 15 different boxes with 10 animals each. The boxes were moved into 3 different incubators (5 boxes each) set at 15, 20 and 25°C respectively and the animals were kept under constant culture regime as described above. When the total of 50 polyps per box was reached, it was maintained constant by manually removing the new clones formed. Every week the number of new clones, dead and spontaneous spawning events where recorded.

### Dry weights

Ten animals from each acclimation temperature (AT) were rinsed quickly in pure ethanol and placed singularly in 1.5ml tubes, previously weighted on an analytical scale. The animals were left dry at 80°C in a ventilated incubator for 4 hours. After complete evaporation of fluids, the animals with the tubes were weighed on the same analytical scale and the dry weight calculated.

### Generation of axenic polyps

In order to reduce the total bacterial load and remove the most of associated bacteria (axenic state), animals belonging to the same clonal line, were treated with an antibiotic (AB) cocktail of ampicillin, neomycin, rifampicin, spectinomycin and streptomycin (50 μg/ml each) in filtered (on 0.2μm filter membrane), autoclaved NM (modified from ^21^). The polyps were incubated in the antibiotic cocktail for two weeks in 50ml Falcon tubes (10 animals each). The medium and the antibiotics were changed every day and the tubes 3 times per week. After the treatment the polyps were incubated for 1 week in sterile NM without antibiotics to let them recover before the recolonization. After the 2 weeks AB treatment, the axenic state was checked by smashing single polyps into 1ml sterile NM and by plating 100μl of the homogenate on marine broth plates, successively incubated for 1 week at 20°C. In addition, we performed a PCR with primers specific for the V1-V2 region of the bacterial 16S rRNA gene (27F and 338R). No CFUs on the plates and a weaker signal in the PCR electrophoretic gel compared with wild-type controls were considered evidences of bacteria reduction and axenic state of the animals.

### Bacteria transplantation

For this experiment, the protocol for conventionalised recolonised *Hydra* polyps was modified from ^21^. For each AT, 100 axenic adult polyps were recolonised with the supernatant of 10 acclimated adult polyps (2 polyps from each acclimated culture box), singularly smashed in 5ml of sterile NM. One ml of supernatant was added into single Falcon tubes, containing 10 axenic animals each and filled with 50ml sterile NM. At the recolonization time, additional animals from the original acclimated cultures (1 polyp/box) were collected for DNA extraction and 16S sequencing. After 24 hours, the medium was exchanged to remove tissue debris and non-associated bacteria. One month after recolonization, the recolonised animals were tested for heat stress tolerance as described above (in 3 rounds of 5 recolonized polyps for each AT). At the time of HS, 15 recolonised polyps for each AT, were sampled for DNA extraction and 16S sequencing.

### Heat stress experiment (HS)

Adult polyps for each AT were placed singularly in 6-well plates and incubated at 40°C for 6 hours (adapted from ^22^). The day after, the number of survivors was recorded and the mortality rate calculated.

### Sexual reproduction induction

Animals separated singularly in 6-well plates, were induced for sexual reproduction via light exposure for 10 h ^17^ and temperature shift to 20°C for the animals acclimated at 15°C, and to 25°C for those acclimated at 20 and 25°C. At each fertilization event, sperm from a single induced male were pipetted directly onto each oocyte pack. Fertilization was performed within 3 hours after spawning. The developing animals were then cultured for 1 month under different temperatures (15, 20 or 25°C).

### Offspring survival test

Ten female polyps from each of the three ATs and one male polyp from the standard culture conditions, were induced separately for spawning. After spawning the adult polyps were removed and the oocyte packs fertilized as described above. Fertilization was confirmed by observation under a binocular of the oocytes first cleavages. After fertilization each oocyte pack was split with a scalpel in 3 parts that were transferred into 3 distinct Petri dishes. The 3 oocyte pack sub-portions were placed into 3 different incubators, set at 15, 20 and 25°C respectively and let develop for one month. Right after fertilization and after one month of development, pictures of the oocytes and the juvenile polyps were acquired for successive counting through ImageJ. Ratios between initial number of oocytes and survived juvenile polyps was calculated and survival rate estimated.

### Bacteria vertical transmission test

Five female polyps from each of the three ATs and one male polyp from the standard conditions, were induced separately for spawning as described above. Immediately after spawning the parental polyps were collected, frozen in liquid N and stored at −80°C for successive DNA and RNA extraction. Five not induced female polyps from each of the three ATs were also collected, frozen and stored for DNA extraction. Oocyte packs were fertilised, split in 3 parts each and let develop for one month at the three different developing temperatures (DTs), as described for the offspring survival test. After one month of development, the juvenile polyps were collected, frozen in liquid N and stored at −80°C. DNA was extracted from both the adults and the offspring as described herein.

### DNA extraction

DNA was extracted from adult polyps starving for 3 days before sacrifice and from never fed juveniles. The recolonized animals were not fed for the whole duration of the AB treatment and the transplantation test (7 weeks in total). Animals were washed two times with 2ml autoclaved MQ, instantly frozen in liquid N without liquid and stored at −80°C until extraction. The gDNA was extracted from whole animals with the DNeasy^®^Blood & Tissue Kit (Qiagen, Hilden, Germany), as described in the manufacturer’s protocol. Elution was done in 50μl and the eluate was stored at −80°C until sequencing. DNA concentration was measured by gel electrophoresis (5μl sample on 1.2% agarose) and by spectrophotometry through Nanodrop 3300 (Thermo Fisher Scientific).

### RNA extraction

Adult animals starved for 3 days before sacrifice. Polyps were washed two times with 2ml autoclaved MQ, instantly frozen in liquid N without liquid and stored at −80°C until extraction. Total RNA was extracted from the body column only, with the AllPrep^®^ DNA/RNA/miRNA Universal Kit (Qiagen, Hilden, Germany), as described in the manufacturer’s protocol. RNA elution was done in 20μl of RNAse-free water and the eluates were stored at −80°C until sequencing. RNA concentration was measured through electrophoresis by loading 1μl of each sample on 1% agarose gel and by spectrophotometry through Nanodrop 3300 (Thermo Fisher Scientific).

### 16S RNA sequencing and analysis

For each sample the hypervariable regions V1 and V2 of bacterial 16S rRNA genes were amplified. The forward primer (5′-**AATGATACGGCGACCACCGAGATCTACAC** XXXXXXXX TATGGTAATTGTAGAGTTTGATCCTGGCTCAG-3′) and reverse primer (5′-**CAAGCAGAAGACGGCATACGAGAT** XXXXXXXX AGTCAGTCAGCCTGCTGCCTCCCGTAGGAGT −3′) contained the Illumina Adaptor (in bold) p5 (forward) and p7 (reverse) ^23^. Both primers contain a unique 8 base index (index; designated as XXXXXXXX) to tag each PCR product. For the PCR, 100 ng of template DNA (measured with Qubit) were added to 25 μl PCR reactions, which were performed using Phusion^®^ Hot Start II DNA Polymerase (Finnzymes, Espoo, Finland). All dilutions were carried out using certified DNA-free PCR water (JT Baker). PCRs were conducted with the following cycling conditions (98 °C—30 s, 30 × [98 °C—9s, 55 °C—60s, 72 °C—90s], 72 °C—10 min) and checked on a 1.5% agarose gel. The concentration of the amplicons was estimated using a Gel Doc TM XR+ System coupled with Image Lab TM Software (BioRad, Hercules, CA USA) with 3 μl of O’GeneRulerTM 100 bp Plus DNA Ladder (Thermo Fisher Scientific, Inc., Waltham, MA, USA) as the internal standard for band intensity measurement. The samples of individual gels were pooled into approximately equimolar subpools as indicated by band intensity and measured with the Qubit dsDNA br Assay Kit (Life Technologies GmbH, Darmstadt, Germany). Subpools were mixed in an equimolar fashion and stored at −20 °C until sequencing. Sequencing was performed on the Illumina MiSeq platform with v3 chemistry (2 × 300 cycle kit) ^24^. The raw data are deposited at the Sequence Read Archive (SRA) and available under the project ID PRJNA742683.

The 16S rRNA gene amplicon sequence analysis was conducted through the QIIME 1.9.0 package ^25^. Sequences with at least 97% identity were grouped into OTUs and clustered against the QIIME reference sequence collection; any reads which did not hit the references, were clustered *de novo*. OTUs with less than 50 reads were removed from the dataset to avoid false positives which rely on the overall error rate of the sequencing method ^26^. Samples with less than 3600 sequences were also removed from the dataset, being considered as outliers. For the successive analysis the number of OTUs per sample was normalized to that of the sample with the lowest number of reads after filtering.

Alpha-diversity was calculated using the Chao1 metric implemented in QIIME. Data were subjected to descriptive analysis, and normality and homogeneity tests as described herein. When normality, homogeneity and absence of significant outliers assumptions were met; statistical significance was tested through one-way ANOVA. When at least one of the assumptions was violated, the non-parametric Kruskal-Wallis test was performed instead. When a significant difference between treatments was stated, post-hoc comparisons were performed in order to infer its direction and size effect.

Beta-diversity was calculated in QIIME according with the different β-diversity metrics available (Binary-Pearson, Bray-Curtis, Pearson, Weighted-Unifrac and Unweighted-Unifrac). Statistical values of clustering were calculated using the nonparametric comparing categories methods Adonis and Anosim.

Bacterial groups associated with specific conditions were identified by LEfSe (http://huttenhower.sph.harvard.edu/galaxy) ^27^. LEfSe uses the non-parametric factorial Kruskal-Wallis sum-rank test to detect features with significant differential abundance, with respect to the biological conditions of interest; subsequently LEfSe uses Linear Discriminant Analysis (LDA) to estimate the effect size of each differentially abundant feature.

### Transcriptome analyses

The analysis was performed on five animals from each AT in two sequencing runs. mRNA sequencing with previous poly-A selection was performed for 15 libraries on the Illumina HiSeq 4000 platform, with 75bp and 150 bp paired-end sequencing respectively. The quality of raw reads was assessed using FastQC v0.11.7 (Andrews, 2014). Trimmomatic v.0.38 ^28^ was then applied to remove adaptors and low-quality bases whose quality scores were less than 20. Reads shorter than 50 bp were removed, and only paired-end reads after trimming were retained. Reads were mapped to the Ensembl metazoa *Nematostella vectensis* genome (release 40) using the splice-aware aligner hisat2 v2.1.0 ^29^ with rna-strandness RF option and default parameters (**Table S1**).

RNA-seq data was used to improve the predicted *N. vectensis* gene model downloaded from Ensembl Metazoa database release 40. Using mapped reads from each temperature condition as input, StringTie v2.0 ^30^ and Scallop v0.10.4 ^31^ were applied to perform genome guided transcriptome assemblies. The assembled transcripts were subsequently compared and merged using TACO ^32^. This produced 42488 genes with 81163 transcripts, among which 21245 genes had significant matches (blastx with parameter e-value 1^e-5^) with proteins in the SwissProt database. Assembled genes were compared with the Ensembl gene model using gffCompare v0.11.2 ^33^, from which genes with lower blastx e-value were selected. Ensembl genes without matching assembled genes were retained, and assembled genes without matching Ensembl genes but with significant matching SwissProt proteins were added to the gene model. The final gene model included 20376 Ensembl genes, 4400 improved genes and 2751 novel assembled genes (**Table S2**). The gene model statistics and the completeness of gene models were assessed using BUSCO v3 ^34^ on the Metazoa dataset that consisted of 978 core genes (**Table S3**).

Total counts of read fragments aligned to the annotated gene regions were derived using FeatureCounts program (Subread-2.0.0) ^35^ with default parameters. Genes whose combined counts from all samples were lower than 5 counts per million (cpm) mapped reads were excluded from the analyses. Differential expression analyses were performed in parallel using DESeq2 (v1.28.1) BioConductor package ^36^, and limma (voom v3.44.3) package ^37^. Differentially expressed genes (DEGs, **Table S4**) were determined based on False Discovery rate (FDR, Benjamini-Hochberg adjusted p-value ≤ 0.05). Gene ontology annotation was derived from the best matching SwissProt proteins. Enriched GO-terms in DEGs were identified by the topGO (v2.40.0) BioConductor package (**Table S5**).

## Results

### Long-term acclimation at high temperature increases heat resistance in *Nematostella vectensis*

Before starting the acclimation experiment, we propagated a single female polyp to 150 clones and split these clones into 15 different cultures with 10 clonal animals each, to ensure the same genotype in all acclimation regimes. We further propagated these animals to 50 animals per culture and constantly maintained this number over the course of the experiment. Subsequently, we aclimated these independent cultures at low (15°C), medium (20°C) and high temperature (25°C) (five cultures each) for the period of 3 years (160 weeks) (**Figure 1**).

**Figure 1.**
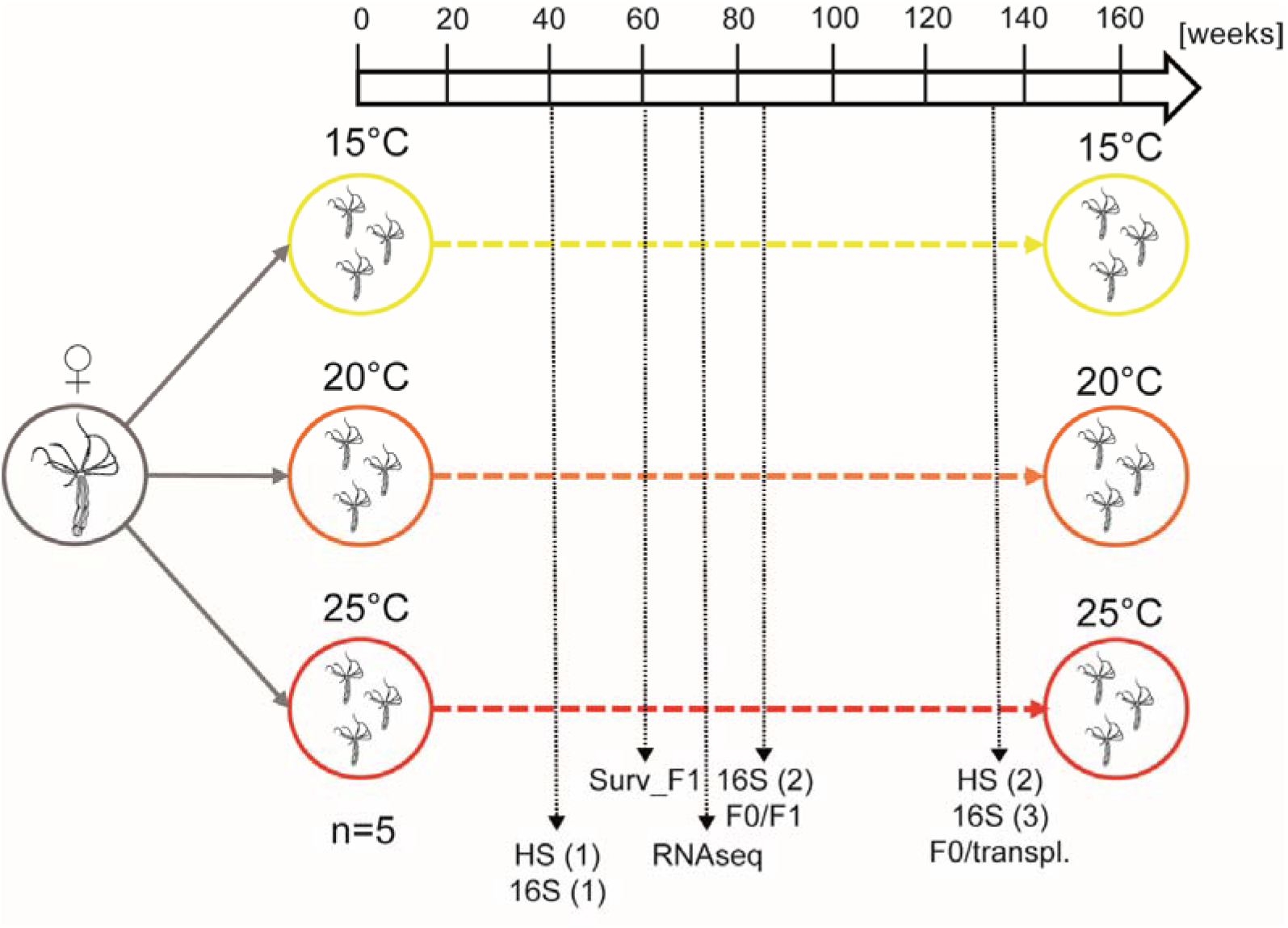
Experimental setup. A single female polyp from the standard culture conditions (16‰ ppt, 20°C) was isolated and propagated via clonal reproduction. When a total of 150 new clones was reached, they were split into 15 different culture boxes of 10 animals each. The boxes were put at three different acclimation temperature (AT) (15, 20 and 25°C, 5 boxes each) and the number of animals/box was kept equal to 50. Heat stress experiments (HS) (6h, 40°C) where performed at 40 and 132 weeks of acclimation (woa). Sexual reproduction was induced at 60 and 84 woa for the juveniles survival test (Surv_F1) and the bacteria vertical transmission test (F0/F1). At 40, 84 and 132 woa samples were collected for 16S sequencing (16S); at 76 woa sampling for RNA sequencing was performed.

After 40 weeks of acclimation (woa), we tested, for the first time, the heat tolerance of acclimated polyps as a proxy for acclimation. We individually incubated polyps of each acclimated culture in ten replicates for 6 hours at 40°C and recorded their mortality (**Figure 2-A**). Already after 40 woa, significant differences in the mortality rates of clonal animals were detectable. While all animals acclimated to low temperature died after the heat stress, animals acclimated at 20°C and 25°C showed a significantly higher survival rate of 70% and 30%, respectively (**Figure 2-A**). We repeated the measurement of heat tolerance two years later (132 woa). Interestingly, we observed a drastic increase in fitness in animals acclimated at high temperature, while the animals acclimated at 15°C and 20°C showed 100% mortality (**Figure 2-A**).

**Figure 2.**
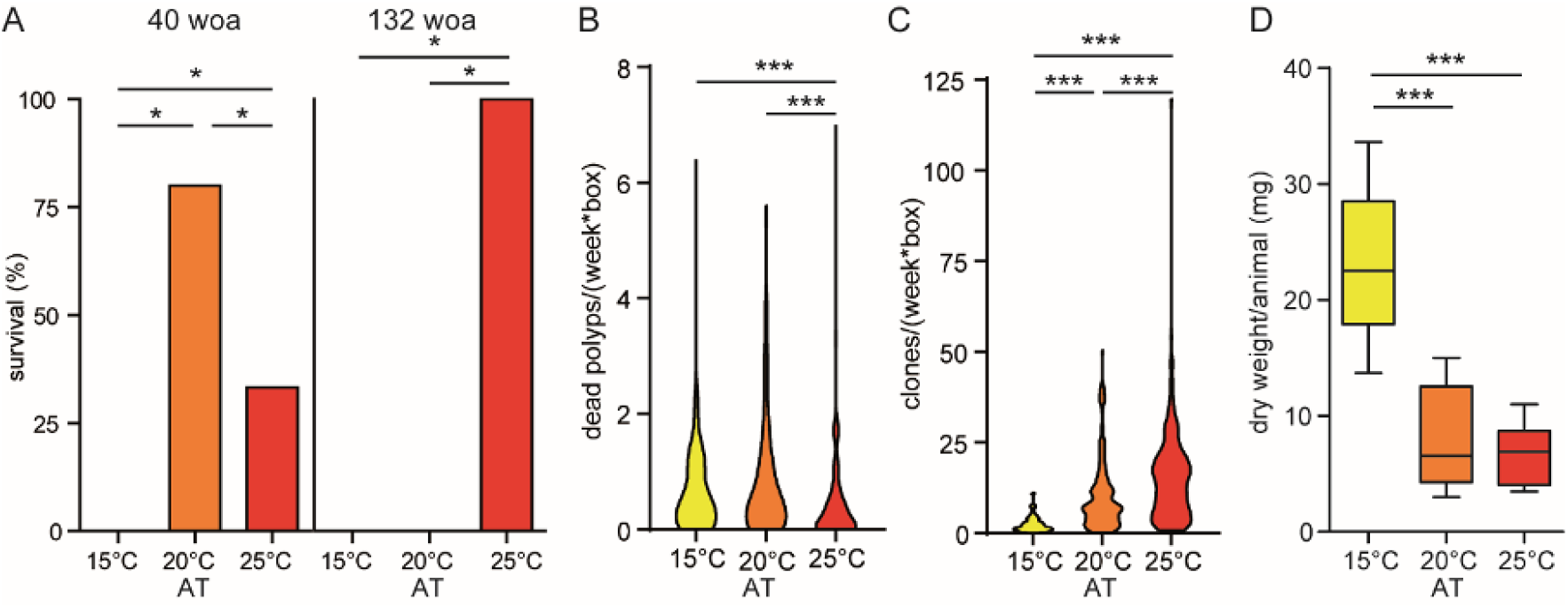
Phenotypic plasticity in response to thermal acclimation. (A) Survival of acclimated polyps after heat stress (40°C, 6 h). Statistical analyses was performed by a Fisher’s exact test (n= 10 (40 woa), n = 5 (132 woa)). (B) Average of dead polyps per week and box over the course of the experiment (E) Average of clones generated per week per 50 animals over the course of the experiment. (D) Dry weights of acclimated polyps at the end of the experiment (170 woa) (n=10): Statistical analyses in B, C and D were performed by one-way ANOVA followed by Tukey’s post-hoc comparisons (* = p ≤ 0.05, ** = p ≤ 0.01, *** = p ≤ 0.001).

We also monitored the mortality rate in the acclimated cultures over the course of the experiment (**Figure 2-B**). While the mortality in cultures acclimated at 15°C and 20°C was below 0.5 polyps per week, the mortality rate at 25°C was significantly reduced in cultures acclimated at 25°C. An additional phenotypic difference between the acclimated animals was the clonal growth, as animals acclimated at 25°C propagated asexually nearly seven times more than animals acclimated at 15°C (**Figure 2-C**). This may explain the differences in body size, where animals acclimated at 15°C were more than three times bigger than the animals acclimated at 20 and 25°C (**Figure 2-D**). The different ATs affected also the fecundity of the animals: the polyps acclimated at the high ATs showed a significantly higher number of spontaneous spawning events recorded along the whole course of the experiment, compared with the 15°C acclimated animals that never spawned if not artificially induced (**Figure S1**).

These results indicate that *N. vectensis* possesses remarkable plasticity at long-term temperature acclimation realized through differences in thermal tolerance, body size, asexual propagation and fecundity. In the following, we analysed the associated microbiota and host transcriptomic responses as a source of thermal acclimation in *N. vectensis*.

### Thermal acclimation leads to dynamic, but reliable changes in the microbiota

To monitor the dynamic changes in the associated microbiotas of acclimated animals, we sampled single polyps from each of the 15 clonal cultures at 40, 84 and 132 woa and compared their associated microbiota by 16S rRNA sequencing (**Figure 1**). To determine the impact of AT and sampling time point on the assemblage of the bacterial community, we performed principal coordinates analysis (PCoA) (**Figure 3-A and B**).

**Figure 3.**
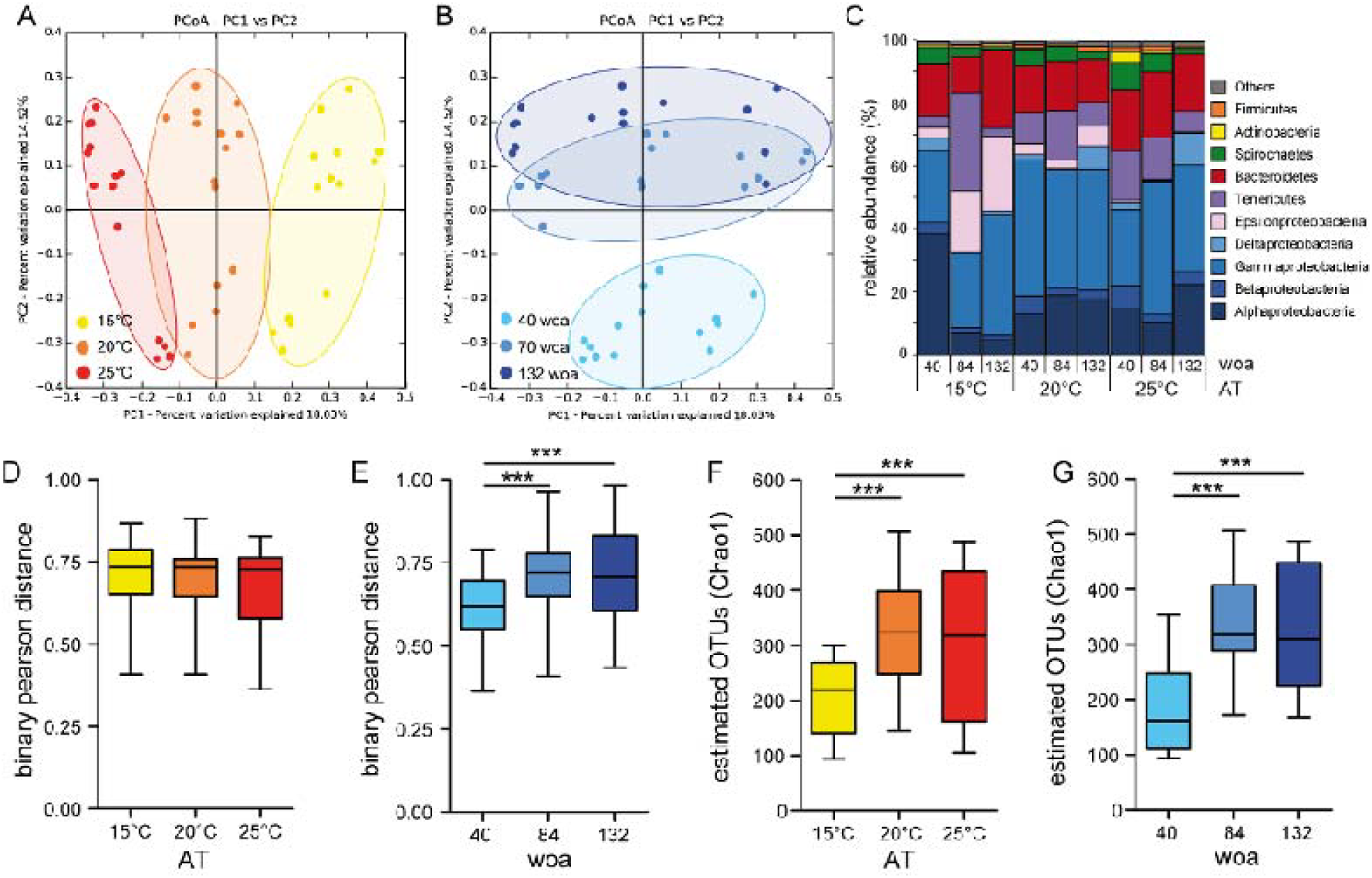
Bacterial community changes in response to thermal acclimation. (A) PCoA (based on binary-pearson metric, sampling depth = 3600) illustrating similarity of bacterial communities based on AT. (B) PCoA (based on pinary-pearson metric, sampling depth = 3600) illustrating similarity of bacterial communities based on woa. (C) Relative abundances of principal bacterial groups, the abundances were summarized under the relative higher taxonomic categories (classes and phyla) and reported as percentages of the total. (D) β-diversity distances within each AT (E) β-diversity distances within woa. Statistical analyses were performed using a non-parametric Kruskal-Wallis test followed by Dunn′s post hoc comparisons (p ≤ 0.01 **, p ≤ 0.001 ***). (F) α-diversity (Chao1) comparison by AT (max rarefaction depth = 3600 (G) α-diversity (Chao1) comparison by woa (max rarefaction depth = 3600), statistical analyses was performed by using one-way ANOVA followed by Tukey’s post hoc comparisons (p ≤ 0.01 **, p ≤ 0.001 ***).

While principal component 1 (PC1) mostly separates samples according to the AT (**Figure 3-A**), PC2 correlates with the different sampling time points (**Figure 3-B**). Using five different β-diversity metrics, we found that bacterial colonization is significantly influenced by both AT and sampling time point (**Table 1**).

**Table 1.** Statistical analysis determining influence of AT and woa on the bacterial colonization. (number of permutations =999).

| parameter | beta-diversity metric | Adonis |  | Anosim |  |
| --- | --- | --- | --- | --- | --- |
|  |  | R<sup>2</sup> | P value | R | P value |
| AT | Binary-Pearson | 0.208 | 0.001 | 0.544 | 0.001 |
|  | Bray-Curtis | 0.219 | 0.001 | 0.466 | 0.001 |
|  | Pearson | 0.256 | 0.001 | 0.360 | 0.001 |
|  | Weighted-Unifrac | 0.147 | 0.001 | 0.238 | 0.001 |
|  | Unweighted-Unifrac | 0.193 | 0.001 | 0.521 | 0.001 |
| woa | Binary-Pearson | 0.230 | 0.001 | 0.608 | 0.001 |
|  | Bray-Curtis | 0.199 | 0.001 | 0.372 | 0.001 |
|  | Pearson | 0.217 | 0.001 | 0.277 | 0.001 |
|  | Weighted-Unifrac | 0.149 | 0.001 | 0.173 | 0.001 |
|  | Unweighted-Unifrac | 0.192 | 0.001 | 0.498 | 0.001 |

Assigning the different microbial communities by the sampling time points revealed a shared clustering after 84 and 132 weeks of acclimation (woa) (**Figure 3-B**), suggesting a stabilization within the microbial communities after around 2 years of acclimation. In contrast, assigning the samples by AT revealed a clear clustering of the microbial communities (**Figure 3-A**) with the bacterial communities acclimated at 20°C clustering between the two extremes (15°C and 25°C). This indicates that the three different ATs induced differentiation of three distinct microbial communities since the beginning of the acclimation process and that this differentiation is more severe between the extreme ATs. While most bacterial groups maintain a stable association with *N. vectensis* (**Figure 3-C**), bacteria that contribute to the differentiation at the end of the acclimation process, are Alphaproteobacteria, that significantly increase at high temperature (Two-way ANOVA, p<0.01) and Epsilonproteobacteria, that significantly increase at low temperature (two-way ANOVA, p<0.001) (**Figure 3-C**).

Using the Binary-Pearson distance matrix, we calculated the distances between samples within all three acclimation regimes (**Figure 3-D**) and sampling time points (**Figure 3-E**). Continuous acclimation under the different temperature regimes revealed no differences in the within-treatment distances (**Figure 3-D**), indicating a similar microbial plasticity at all three ATs. In contrast, Binary-Pearson distances of the different sampling time points significantly increased between 40 and 84 woa (**Figure 3-E**) and stabilized between 84 and 132 woa. Interestingly, the α-diversity of bacteria associated with acclimated polyps was significantly higher at 20 and 25°C, compared to those associated with polyps acclimated at 15°C (**Figure 3-F**). As for the β-diversity, the α-diversity was significantly increasing within the first 84 woa and stabilized between 84 and 132 woa (**Figure 3-G**).

Altogether, these results show that the microbiota of *N. vectensis* reacts plastically to environmental changes. The microbial composition changes stabilize within two years of acclimation indicating a new homeostatic bacterial colonization status.

### Thermal acclimation induces a robust tuning of host transcriptomic profiles

To evaluate the contribution of host transcriptional changes to the observed increased thermal tolerance in animals acclimated at high temperature, we analysed gene expression profiles of *N. vectensis* after 75 woa (**Figure 1**). We sampled from each replicate culture one animal, extracted its mRNA and sequenced it by Illumina HiSeq 4000. The constant acclimation at 15, 20 and 25°C induced a robust tuning of the host transcriptomic profiles (**Figure 4-A**).

**Figure 4.**
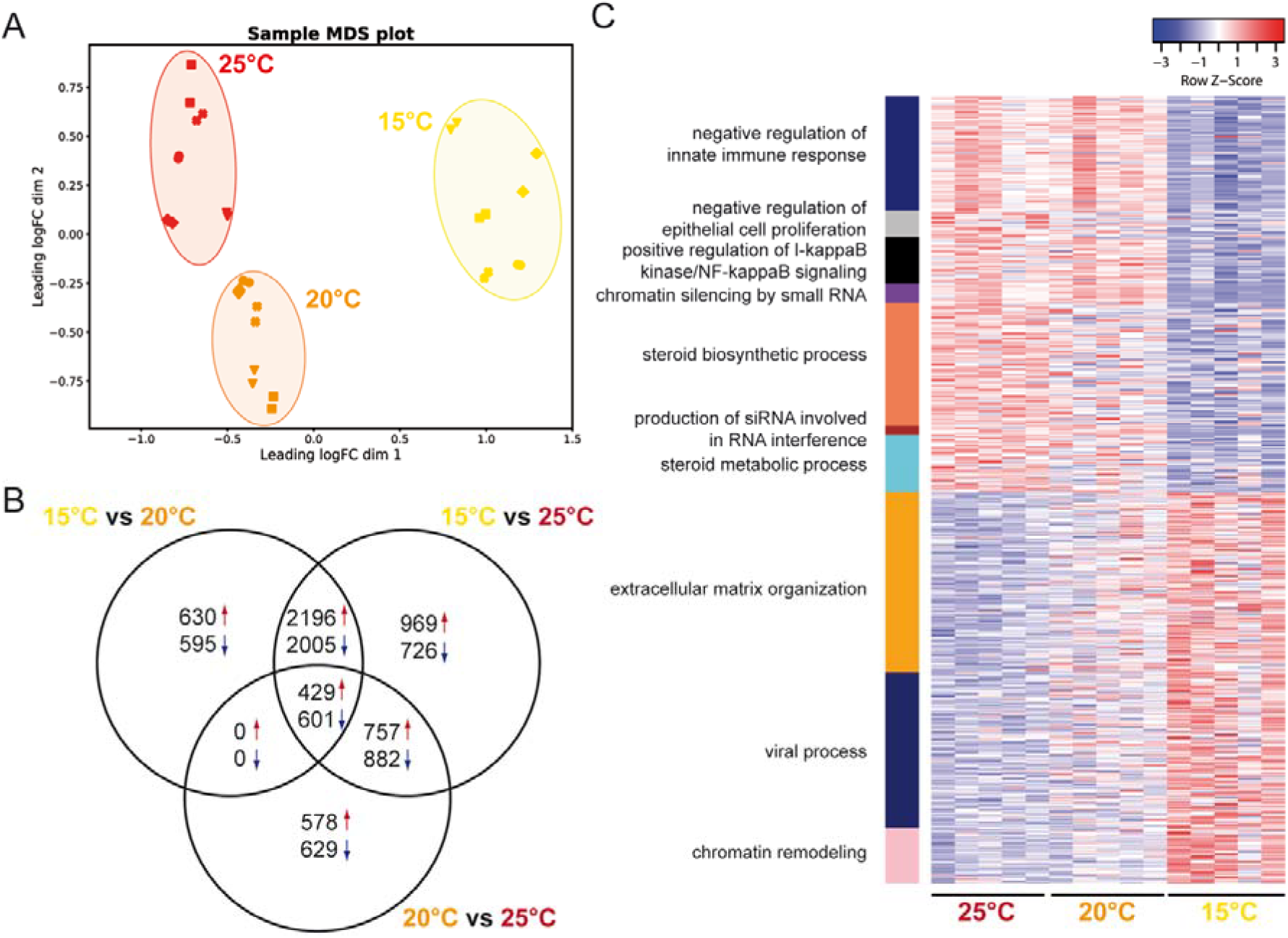
Host transcriptome changes after thermal acclimation. (A) MDS plot showing the clustering of the transcriptome samples according to the AT of the acclimated animals (samples were sequenced in technical replicates, indicated by the different symbols) (B) Venn diagram showing the differentially expressed genes within the three ATs pairwise comparisons. (C) Heat-map of differentially expressed genes in enriched GO term categories significantly enriched in the comparison between 15 and 25°C acclimated polyps.

In pairwise comparisons, we determined the differentially expressed (DE) genes (**Figure 4-B**) in all acclimated animals. While the comparison of transcriptomic profiles from polyps acclimated at 15 and 25°C revealed the highest number of DE genes, the comparison of 20 and 25°C acclimated animals revealed the lowest number of DE genes. In all three comparisons, we observed a similar fraction of up- and down-regulated DE genes (**Figure 4-B**).

To retrieve potential molecular processes and signalling pathways enriched at the different ATs, we performed a gene ontology (GO) enrichment analysis and concentrated on GO categories significantly enriched in the comparison between 15 and 25°C acclimated polyps (**Figure 4-C, Table S3**). Animals acclimated to high temperature significantly increased expression in genes involved in innate immunity, gene regulation, epithelial cells proliferation, steroid biosynthesis and metabolism (**Figure 4-C, Table S5**). While genes associated with enriched GO categories show opposite expression levels at 15°C and 25°C, an intermediate expression level was evident in the animals acclimated at 20°C (**Figure 4-C**). The animals acclimated to low temperature showed upregulation of genes associated with viral processes, which seems to be compatible with their general lower viability.

### Transplantation of acclimated microbiota induces differences in heat tolerance

To disentangle the effects of transcriptomic and bacterial adjustments on thermal tolerance of acclimated polyps, we performed microbial transplantation experiments. We generated axenic non-acclimated animals and recolonized these animals with the microbiota of long-term acclimated polyps from the same clonal line. We smashed acclimated animals and used these suspensions, containing the acclimated microbiota, for the recolonization of axenic animals. We maintained microbiota-transplanted animals for one month at 20°C to allow the adjustment of a stable colonization.

To evaluate the success of bacteria transplantation, we performed 16S rRNA gene sequencing of 45 recolonized animals. PCoA analysis revealed that the transplanted microbiota cluster according to the acclimated source microbiota one month after transplantation (**Figure 5-A**, **Table 2**).

**Figure 5.**
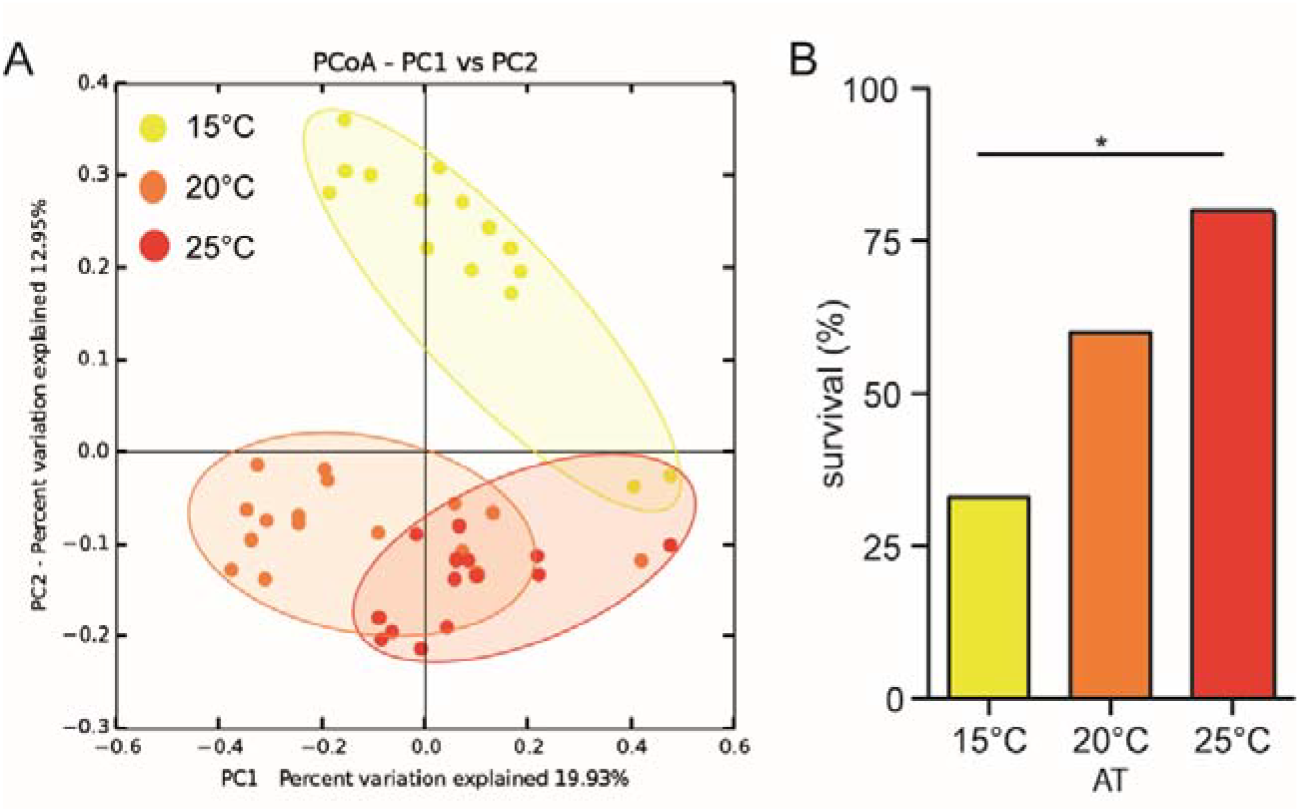
Transplantation of acclimated microbiota confers thermal resistance. (A) PCoA (based on binary-pearson metric, sampling depth = 3600) illustrating similarity of transplanted bacterial communities based on AT of source microbiota (B) Heat stress survival of polyps recolonized with microbiota of acclimated animals. Statistical analyses were performed by pairwise Fisher’s exact test (n = 15, * p = 0.025.

**Table 2.** Statistical analysis determining influence of AT of source microbiota on bacterial colonization. (number of permutations = 999).

| parameter | beta-diversity metric | Adonis |  | Anosim |  |
| --- | --- | --- | --- | --- | --- |
|  |  | R<sup>2</sup> | P value | R | P value |
| AT of source microbiota | Binary-Pearson | 0.199 | 0.001 | 0.486 | 0.001 |
|  | Bray-Curtis | 0.183 | 0.001 | 0.346 | 0.001 |
|  | Pearson | 0.165 | 0.001 | 0.194 | 0.001 |
|  | Weighted-Unifrac | 0.161 | 0.001 | 0.272 | 0.001 |
|  | Unweighted-Unifrac | 0.184 | 0.001 | 0.416 | 0.001 |

Subsequently we tested the microbiota-transplanted animals for their heat tolerance as previously performed for the acclimated animals. The recolonized animals showed clear differences in mortality depending on the microbial source used for transplantation. A significant gradient in survival was evident from the animals recolonized with the 15°C-acclimated microbiota (33%) to those recolonized with the 25°C-acclimated microbiota (80%) (**Figure 5-B**). The animals transplanted with the 20°C-acclimated microbiota showed an intermediate survival (60%).

These results indicate that the high thermal tolerance of animals acclimated to high temperature can be transferred to non-acclimated animals by microbiota transplantation alone. Therefore, we conclude, that microbiota-mediated plasticity provides a rapid mechanism for a metaorganism to cope with environmental changes.

Through the LEfSe analysis, we were able to detect bacterial OTUs differentially represented between the polyps acclimated at 15 and 25°C, and in the corresponding transplanted animals (**Table S6**). These bacteria belong to the families Phycisphaeraceae, Flavobacteriaceae, Emcibacteraceae, Rhodobacteraceae, Methylophilaceae, Francisellaceae, Oceanospirillaceae and Vibrionaceae, which are known to include various commensals, symbionts and pathogens of marine organisms. Therefore, the OTUs overrepresented in the 25°C microbiota may constitute good candidates for providing thermal resistance to their host.

### Acclimated microbiota and thermal tolerance are transmitted to next generation

In a next step, we tested if the acclimated microbiota influencing adults’ thermal tolerance is also affecting thermal tolerance of the offspring. Therefore, ten female polyps from each long-term acclimated culture and one non-acclimated male polyp were induced separately for spawning. All oocyte packs were fertilized with the sperm of the same male polyp, split into three parts, counted and let develop for one month at the three different temperatures (developing temperature - DT) in a full factorial design (**Figure 6-A**).

**Figure 6.**
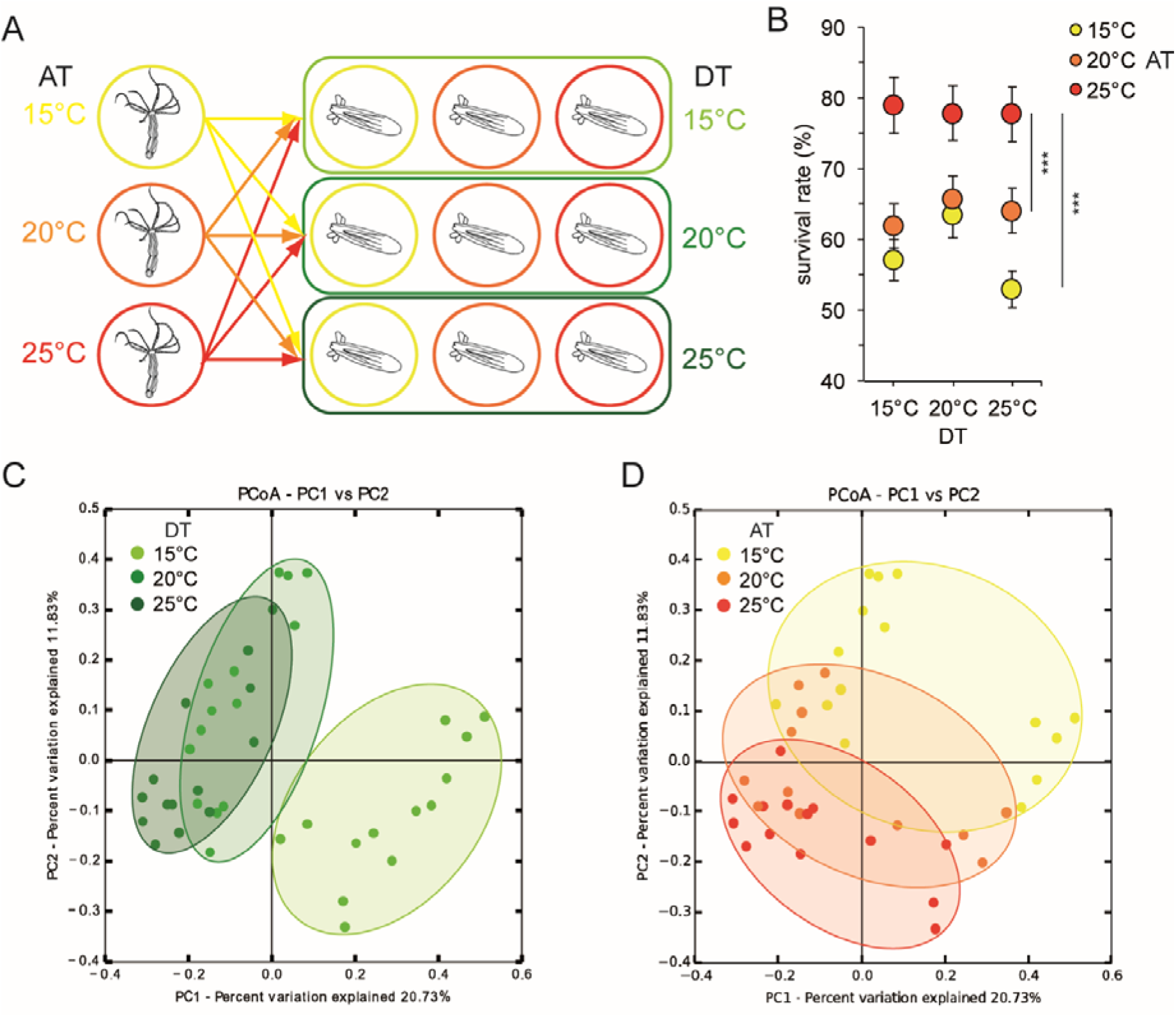
Transmission of thermal tolerance to the offspring. (A) Experimental scheme: acclimated females from each AT were induced for sexual reproduction. All oocyte-packs were fertilized with the sperms from a single male polyp from the standard conditions. After fertilization, each embryo-pack was split in 3 parts and placed at different DT (15, 20 or 25°C). After one month of development, survival rate and bacterial colonisation were analysed. (B) Offspring survival rate (ratio between initial number of oocytes and survived juveniles polyps were calculated), a Kruskal-Wallis test was performed followed by Dunn′s post-hoc comparisons (n=10; *** p ≤ 0.001). (C) PCoA (based on Binary-Pearson metric, sampling depth = 24500) illustrating similarity of bacterial communities according with offspring DT, (D) PCoA (based on Binary-Pearson metric, sampling depth = 24500) illustrating similarity of bacterial communities according with mothers’ AT.

After one month of development, the survived juvenile polyps were counted and corresponding survival rates were calculated (**Figure 6-B**). The offspring from the mothers acclimated at 25°C showed a significant higher overall survival rate compared to the offspring from polyps acclimated at medium and low temperature. In contrast, the offspring of polyps acclimated at 15°C showed the lowest survival rate at 25°C DT (**Figure 6-B**). In a second step, the juvenile polyps were subjected to 16S rRNA sequencing to evaluate the transmission of acclimated microbes to the next generation. PCoA revealed a significant clustering of samples according to both DT of the juveniles and AT of mother polyps (**Figure 6-C and D, Table 3**). While, on average, around 50% of bacterial variation can be explained by the DT of the juvenile polyps, around 20% of the bacterial colonization in juveniles can be explained by the acclimation temperature of the mother polyps (**Table 3**).

**Table 3.** Statistical analysis determining influence of mothers’ AT and offspring DT on bacterial colonization of F1 generation polyps. (number of permutations = 999).

| parameter | beta-diversity metric | Adonis |  | Anosim |  |
| --- | --- | --- | --- | --- | --- |
|  |  | *R*<sup>2</sup> | *P* value | *R* | *P* value |
| **AT of mother polyps** | Binary-Pearson | 0.170 | 0.001 | 0.373 | 0.001 |
|  | Bray-Curtis | 0.133 | 0.001 | 0.237 | 0.001 |
|  | Pearson | 0.139 | 0.001 | 0.166 | 0.002 |
|  | Weighted-Unifrac | 0.093 | 0.024 | 0.091 | 0.010 |
|  | Unweighted-Unifrac | 0.116 | 0.001 | 0.148 | 0.005 |
| **DT** | Binary-Pearson | 0.262 | 0.001 | 0.696 | 0.001 |
|  | Bray-Curtis | 0.260 | 0.001 | 0.621 | 0.001 |
|  | Pearson | 0.338 | 0.001 | 0.542 | 0.001 |
|  | Weighted-Unifrac | 0.300 | 0.001 | 0.408 | 0.001 |
|  | Unweighted-Unifrac | 0.211 | 0.001 | 0.413 | 0.001 |

These results demonstrate that the acquired thermal tolerance in the maternal generation is transmitted to the next generation. The fact that also parts of the acclimated microbiota is transmitted and persisting in the juvenile F1 polyps suggests that vertically transmitted acclimated bacteria can be adaptive to high temperature.

## Discussion

### Long-term acclimation promotes heat tolerance in *N. vectensis*

The ability for marine animals to adapt to future thermal scenarios is of pivotal importance for the maintenance of biodiversity and ecosystem functioning. Recent studies indicate that sessile marine animals, like corals, sponges or anemones, could adapt more rapidly than expected to climate change ^2,38–42^. Recent and long-term observations in the field displayed higher heat tolerance of corals pre-exposed to thermal stress compared to unexposed ones and showed that wild populations are slowly becoming less sensitive than they were in the past ^43–45^. In our study, the host’s thermal resistance showed an increase along with the acclimation time. It is important to point out that the standard culture temperature for *N. vectensis* in the lab is 20°C. The animals maintained at 20°C, therefore, have been acclimated to this condition for a long time and this might explain their highest survival at 40 woa. Interestingly, the animals acclimated at 15°C showed at both time points 100% mortality, indicating that these animals would not be able to survive extreme temperature events. Our results are consistent with other studies that investigated the acclimation capacity of corals in lab experiments. Pre-acclimated individuals of *Acropora pruinosa*, a scleractinian coral, did not bleach when exposed to successive heat stress ^42^. Also in the field, *Acropora hyacinthus* showed less mortality after heat stress when acclimated to wide temperature fluctuations, than when acclimated to less variable environments ^46^. These different resistances are correlated to an adaptive plasticity in the expression of environmental stress response (ESR) genes ^47^ and the presence of an advantageous microbiota ^48^, but a causative relation was not shown in both cases. In our study, we disentangled the contribution of host gene expression and microbiota to temperature acclimation in cnidarian, by the use of microbial transplantation experiments in a single host genotype background.

### Microbiota plasticity promotes metaorganism acclimation

Shifts in the composition of bacterial communities associated with marine animals in response to changes in environmental factors (i.e. temperature, salinity, pH, light exposure, oxygen and CO2 concentrations, etc.), has been demonstrated in numerous studies ^18,49–55^. In some cases these changes in microbiota composition correlated with a higher fitness of acclimated animals ^53^, but causal connections are rare. An experimental replacement of a single bacterium and subsequent demonstration of acquired heat tolerance by the host, was only shown in aphids ^56^. To infer if and to what extent the acclimated microbiota confers thermal resistance, we performed transplantation experiments of microbiotas from acclimated animals to non-acclimated ones. These experiments proved that polyps transplanted with the microbiota from animals acclimated at 25°C, acquired a higher thermal tolerance than those transplanted with the 15°C acclimated microbiota. It is important to point out that the animals selected as receivers for this experiment were all clones of the same age, size and shared the same life history, since they came from the same culture box and belonged to the same clonal line as the acclimated donors. With this experimental setup, we were able to disentangle host and microbiota contribution to thermal acclimation and proved that acclimated bacteria can act as heat tolerance promoting bacteria (HTPB).

The acclimation of the microbial community is a highly dynamic process that started within the first weeks after environmental shift, and most of the bacterial β-diversity adjustments happened until 84 woa. Afterwards the microbial community likely reached a stable and homeostatic state. Previous studies on corals ^57,58^ detected the presence of a “core microbiota”, defined as a cluster of microbial species that are persistent either temporally and/or among different environments or locations, are associated with host-constructed niches, and therefore less sensitive to changes in the surrounding environment. Members of the core microbiota may not necessarily represent the most abundant groups of the community but are hypothesized to exert pivotal functions for the maintenance of the holobiont homeostasis. In contrast, a “dynamic microbiota” exists that varies depending on species, habitat and life stage and is likely a product of stochastic events or a response to changing environmental conditions ^58^. Also in *N. vectensis* it seems that during the acclimation process, a core microbiota remained stable in all acclimated polyps, while a more dynamic part of the microbiota changed by either increasing or decreasing in punctual abundances.

The increase in α-diversity indicates either the acquisition of new bacterial species from the surrounding or a higher evenness in species abundances, where OTUs that were rare at the beginning of the experiment and at lower temperature, became more abundant and therefore detectable. The acquisition of new bacterial species during lab experiments appears unlikely since the polyps are isolated from their natural environment. Nevertheless, the acclimated animals are not maintained under sterile conditions and thus an exchange of microbial species with the culture medium and from the food supply cannot be excluded. As already pointed out in numerous studies ^59–61^, higher microbial diversity enhances the ability of the host to respond to environmental stress by providing additional genetic variability, and corals exposed to heat stress exhibit increased microbiota β-diversity ^62^.

In addition to the changes in species composition and relative abundances, the associated microbial species can evolve much more rapidly than their multicellular host ^8^. Rapidly dividing microbes are predicted to undergo adaptive evolution within weeks to months ^63^. Therefore, adaptation of the host can also occur via symbiont acquisition of novel genes ^64^, via mutation and/or horizontal gene transfer (HGT) ^8^. Therefore, it is possible that, even if a certain bacterial species didn’t significantly change in abundance between the different ATs, it may have acquired new functions and adapted to the new conditions within the course of the experiment.

Alphaproteobacteria and Gammaproteobacteria constitute main microbial colonizers of corals ^57,65^ and of *N. vectensis* ^18,66^. The increased thermal tolerance of animals acclimated at high temperatures is often associated with an increase in abundance of these bacterial classes in the associated microbiota ^67,68^. In thermally stressed animals, Alphaproteobacteria constitute an important antioxidant army within the coral holobiont ^69^ and together with members of the Gammaproteobacteria class, were found to significantly inhibit the growth of coral pathogens (e.g. *V. coralliilyticus* and *V. shiloi*) ^7,70^. They are also known to exert nitrogen fixation in endosymbiosis with marine animals, providing the host with additional nutrient supply ^71–73^. In our study, Alphaproteobacteria significantly increased in abundance in the animals acclimated at high temperature and most of the bacterial OTUs significantly overrepresented in the animals transplanted with the 25°C acclimated microbiota, belong to the Alpha- and Gammaproteobacteria classes. Among these OTUs, those that could be classified with high confidence, are members of the genera *Sulfitobacter, Francisella* and *Vibrio*, and one Flavobacteriia OTU of the genera *Muricauda*. All these bacterial groups are known to comprise pathogens and symbionts of multicellular organisms ^74^. In particular, *Sulfitobacter* is an endosymbiont of vestimentiferans inhabiting hydrothermal vents, where it performs sulfite oxidation ^75^; *Francisella* is an intracellular pathogen of mammals and various invertebrates and it is supposedly capable of ROS scavenging ^76,77^. Members of the Flavobacteriaceae family are key players in biotransformation and nutrient recycling processes in the marine environment, known intracellular symbionts of insects and intracellular parasites of amoebae ^78^. All these characteristics make them good candidates for providing thermal tolerance to the host.

### Changes in host gene expression may confer acclimation

Previous studies on *Hydra* showed that the cnidarian innate immune system actively controls the composition and the homeostasis of the associated microbiota, and that such associations are both species-specific and life-stage specific ^79–82^. In corals, it has been shown that unacclimated individuals expressed stronger immune and cellular apoptotic responses than acclimated ones, and disease-related metabolic pathways were significantly enhanced in the former ^42^. Moreover, the immune system is sensitive to environmental change ^59^ and colonization by beneficial symbionts might lead to the suppression of the host immune response ^55^. Elements of the innate immune system, including several members of the interleukin signaling cascades and the transcription factor NF-kB, have been characterized in *N. vectensis* and are hypothesised to play similar roles as their vertebrata homologs ^15,83–85^. We hypothesise that the lower expression of genes involved in the innate immune response, plus a positive regulation of the NF-kB signaling observed in the animals acclimated to the higher ATs, indicates a general suppression of the host’s immune response. Animals challenged by unfavourable environmental conditions (high temperature in this case), may suppress their immune reaction to favour the establishment of new symbionts. Interestingly, a GO term comprising genes implicated in viral processes, were also upregulated in the animals acclimated at 15°C, suggesting a possible higher susceptibility of these animals to infections and a possible implication to their lower viability.

On the other hand, steroids and secosteroids from gorgonian and soft corals, have been shown to have antimicrobial and antifouling activity ^86,87^ and ^88^ found in, *N. vectensis*, homologs of genes involved in steroids metabolism in other animals. The upregulation of genes involved in steroid biosynthesis and metabolism in the high ATs animals may indicate a role in chemical defence against pathogens. In addition, it might hint to the contribution of steroid signalling in the regulation of phenotypic plasticity^89,90^, e.g. in body size regulation and reproduction rate in response to different temperatures.

The enhanced production of small RNAs (sRNAs) in the high ATs acclimated animals, and the high regulation of processes involved in chromatin remodelling in the 15°C acclimated animals, suggest a general high gene transcription and translation regulations at these two extreme, not optimal, conditions. Chromatin remodelling processes are implicated in epigenetic modifications and thus possibly inheritable by the offspring ^91^. A recent publications ^92^ analyzed coral-associated bacteria proteomes and detected potential host epigenome-modifying proteins in the coral microbiota. This, in concert with specific symbionts inheritance, may constitute an additional mechanism for thermal resistance transmission along generations and may explain the significantly higher viability of the 25°C acclimated animals’ offspring.

### Acquired thermal tolerance is transmitted to the next generation

The capacity of a species to survive and adapt to unfavourable environmental conditions does not only rely on the adaptability of the adults but also on the survival of the early life stages. Even if the adults are able to acclimate to periodic heat waves and seasonal temperature increases, their offspring may have a much narrower tolerance range ^22,93–95^. It is evident that offspring of marine species, including fishes, mussels, echinoderms and corals can acclimatize to warming and acidifying oceans via transgenerational plasticity (TGP) ^96–103^. Both transmission of epigenetic modifications ^101,104–108^ and microbiota-mediated transgenerational acclimatization (MMTA) ^8,40^ may be involved in the process.

Recently, it was shown in *N. vectensis* that animals acclimated to high temperature transmit thermal resistance to their offspring ^109^. In our experiments, we moved a step forward by exploring the contribution that the microbiota may have in the inheritability of this plasticity. We fertilized oocytes of acclimated females with sperm of a single male in order to keep the genetic variability as low as possible, and cultured the offspring in a full factorial design at 15°C, 20°C and 25°C. As expected, offspring originated from mothers acclimated at 25°C showed the highest survival rate. These results confirmed that polyps acclimated to high temperatures, transmit their acquired thermal tolerance to their offspring, increasing their fitness at high temperature. The fact that offspring from genetically identical mothers show differences in survival rate, indicates either (i) the vertical transmission of HTPB or (ii) the transmission of epigenetic modifications.

For many marine invertebrates, vertical transmission of microbial symbionts are assumed ^110–113^. In particular, species that undertake internal fertilization and brood larvae, tend to preferably transmit their symbionts vertically, whereas broadcast spawners and species that rely on external fertilization are thought to mainly acquire their symbionts horizontally ^114–116^. Bacteria may also be transmitted to the gametes by incorporation into the mucus that surrounds oocyte and sperm bundles ^117–119^. Alternatively, the gametes may acquire bacteria immediately after release by horizontal transmission through water, which contains bacteria released by the parents ^55^. A recent publication showed that *N. vectensis* adopts a mixed mode of symbionts transmission to the next generation, consisting of a differential vertical transmission from male and female parent polyps, plus an horizontal acquisition from the surrounding medium during development ^19^. Consistently, the results of this study suggest the vertical transmission of HTPB.

### Acclimated microbiota-a source for assisted evolution

Microbial engineering (ME) is nowadays regularly applied to agriculture and medicine to improve crop yields and human health ^120^. Pioneering theoretical works, including the Coral Probiotic Hypothesis ^7^ and the Beneficial Microorganisms for Corals (BMC) concept ^121^, suggested that artificial selection on the microbiota could improve host fitness over time frames short enough to cope with the actual and future rates of climate changes. Some studies have started ME on corals as a restoration/conservation option for coral reefs subjected to environmental stresses ^122–124^. Recently, was showed that corals subjected to experimental warming, inoculated with consortia of potentially beneficial bacteria, bleached less compared to corals that received no probiotics ^125^. It needs to be pointed out that MMTA is of pivotal interest because it would be a suitable target for manipulations in perspective of future assisted evolution (AE) programs ^40,125^.

In this study we proved that long-term acclimation induces enormous changes in the physiology, ecology and even morphology of genetically identical animals; that animals exposed to high (sublethal) temperatures can acclimate and resist to heat stress and that this resistance can be transmitted to the next generations and to non-acclimated animals by microbiota transplantation. We have been able to detect specific bacterial groups that could be responsible for providing different thermal tolerances to their host and that may represent good candidates for future assisted-evolution experiments.

## Acknowledgments

This work was supported by the Human Frontier Science Program (Young Investigators' Grant RGY0079/2016 and the DFG CRC grant 1182 “Origin and Function of Metaorganisms” (Project B1).

## Conflict of Interest statement

The authors declare no conflict of interest.

Supporting information file is available online.

## Notes

### Competing Interest Statement

The authors have declared no competing interest.

