## Supplementary Figures for "Microbiota mediated plasticity promotes thermal adaptation in *Nematostella vectensis*"

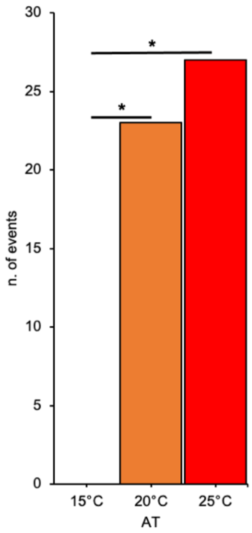


**Figure S1. Number of spontaneous spawning events at each AT along the whole duration of the acclimation experiment.** The spawning events were recorded when egg packs were found in all the boxes (n = 5) from each AT. Differences were tested through Fisher's Exact test (* = $p \leq 0.05$).

**Table S6. Bacterial OTUs overrepresented at 15 and 25°C in both acclimated and recolonized animals**. Through LEfSe were detected the OTUs significantly overrepresented at 15 vs 25°C for both the acclimated animals and the transplanted ones (factorial Kruskal-Wallis test α-value = 0.05, logarithmic LDA score threshold = 2). In the table are reported the OTUs shared between the acclimated and the transplanted animals according with the AT (in grey the classifications that don't reach 80% confidence similarity with reference sequences available in public databases).

| Phylum | Class | Order | Family | Genus | OTU | AT |
|---|---|---|---|---|---|---|
| Proteobacteria [100%] | Betaproteobacteria [100%] | Burkholderiales [100%] | Alcaligenaceae [100%] | *Castellaniella* [100%] | OTU 509480 | 15°C |
| Proteobacteria [100%] | Gammaproteobacteria [100%] | Alteromonadales [96%] | Colwelliaceae [96%] | *Colwellia* [86%] | OTU 349769 | |
| Proteobacteria [100%] | Gammaproteobacteria [100%] | Cellvibrionales [100%] | Spongiibacteraceae [98%] | *Marortus* [87%] | OTU 328 | |
| Planctomycetes [100%] | Phycisphaerae [100%] | Phycisphaerales [100%] | Phycisphaeraceae [100%] | *Algisphaera* [74%] | OTU 135 | 25°C |
| Bacteroidetes [100%] | Flavobacteriia [100%] | Flavobacteriales [100%] | Flavobacteriaceae [100%] | *Muricauda* [98%] | OTU 129 | |
| Proteobacteria [99%] | Alphaproteobacteria [97%] | Emcibacterales [69%] | Emcibacteraceae [69%] | *Emcibacter* [69%] | OTU 42 | |
| Proteobacteria [100%] | Alphaproteobacteria [100%] | Rhodobacterales [100%] | Rhodobacteraceae [100%] | *Sulfitobacter* [98%] | OTU 650063 | |
| Proteobacteria [100%] | Alphaproteobacteria [100%] | Rhodobacterales [100%] | Rhodobacteraceae [100%] | *Sulfitobacter* [58%] | OTU 162 | |
| Proteobacteria [100%] | Betaproteobacteria [100%] | Nitrosomonadales [89%] | Methylophilaceae [83%] | *Methylotenera* [60%] | OTU 232 | |
| Proteobacteria [100%] | Gammaproteobacteria [100%] | Thiotrichales [100%] | Francisellaceae [100%] | *Francisella* [100%] | OTU 144057 | |
| Proteobacteria [100%] | Gammaproteobacteria [100%] | Thiotrichales [100%] | Francisellaceae [100%] | *Francisella* [100%] | OTU 218 | |
| Proteobacteria [100%] | Gammaproteobacteria [100%] | Oceanospirillales [100%] | Oceanospirillaceae [100%] | *Neptunomonas* [78%] | OTU 275 | |
| Proteobacteria [100%] | Gammaproteobacteria [100%] | Vibrionales [100%] | Vibrionaceae [100%] | *Vibrio* [100%] | OTU 939811 | |
